# Independent Associations of Sleep and Walking with Cognition in Parkinson’s Disease: A Perivascular Spaces Analysis

**DOI:** 10.1101/2025.05.16.654573

**Authors:** Francesca Sibilia, Giuseppe Barisano, Maryam Fotouhi, Philipp A. Loehrer, Katherine Wong, Wendy J. Mack, Mark F. Lew, Haidar Dafsari, Jeiran Choupan

## Abstract

Perivascular spaces (PVSs) support brain waste clearance and are influenced by sleep. In Parkinson’s disease (PD), sleep is often disrupted, and can co-occur with gait dysfunction. This study investigated how sleep disruption, walking difficulties, and cognition relate in PD, considering PVS as a mediator. Our cohort included 348 participants -healthy controls (HC, n=59), prodromal (n=117), and PD patients (n=172)-in the Parkinson’s Progression Markers Initiative (PPMI) study. Insomnia and walking issues were assessed by questionnaire, cognition was measured with the Symbol Digit Modalities (SDM) test, and PVS volume fraction in white matter (WM-PVS) and basal ganglia (BG-PVS) was quantified. PVS did not mediate the relationship between sleep or motor problems, on cognition. Walking difficulties were directly linked to poorer cognition, and insomnia to walking deficits. BG-PVS was negatively associated with cognition. PVS burden predicted cognitive function after five years, but not disease progression as indexed by cerebro-spinal fluid (CSF) biomarkers. These findings suggest walking deficits directly relate with cognition in PD, but the relationship is not mediated by PVS.

## 1. Introduction

Parkinson’s disease (PD) is a neurodegenerative disorder characterized by motor and non-motor symptoms^1^. Poor sleep^2^ and walking difficulties^3^ are among the first symptoms of PD, often occurring prior to clinical diagnosis. Increased walking difficulties limit patients’ ability to engage in beneficial physical activity. Physical limitations and sleep disturbances significantly affect cognitive performance in PD: slower walking speed is linked to cognitive decline in older adults with PD^4^. Reduced physical activity is also associated with poorer sleep patterns^5^, and 60–80% of PD patients experience sleep problems such as insomnia, daytime sleepiness, restless legs syndrome, or rapid eye movement sleep behavior disorder (RBD)^6^. Global cognition was found to be associated with balance problems and walking functions, and to be worsened by chronic sleep disruptions in PD.

Sleep is important for the optimal functioning of the glymphatic system, that is responsible for the brain waste clearance^7^. Poor sleep quality can exacerbate neurotoxins and waste molecules accumulation, altering the glymphatic system normal functioning^7^. Perivascular spaces (PVS), an important component of the glymphatic system, are fluid-filled spaces surrounding blood vessels and serve as a conduit for waste clearance and interstitial fluid dynamics^8^. Previous clinical studies highlighted the role of enlarged PVS in white matter in PD^9^ and the association of sleep problems with CSF biomarkers of neurodegeneration^10^. Elevated CSF levels of alpha-synuclein and total Tau (t-Tau) have been associated with increased sleep fragmentation and RBD^11^. Higher levels of PET-measured beta-amyloid have also been linked to greater sleep disturbances in people with RBD, suggesting an association between sleep and abnormal brain protein accumulation^12^. Despite the individual recognition of PVS alterations^9^, sleep disturbances^2^, and walking abnormalities in PD^4^, the interplay among these factors remains underexplored. Understanding how sleep and walking problems may affect PVS integrity and cognition could provide valuable insights into the underlying PD pathophysiology (**Figure1**)^13^.

**Figure1:**
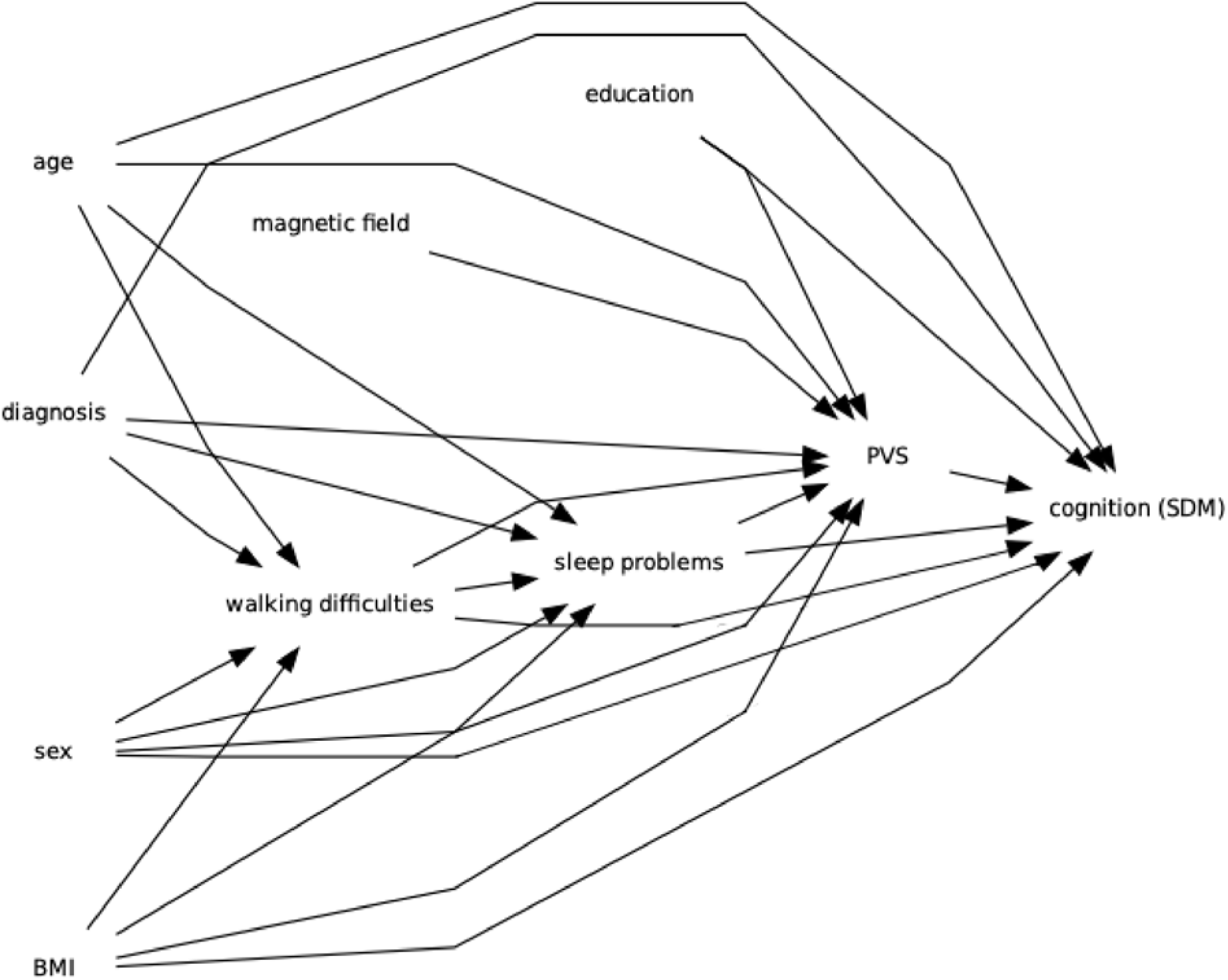
Directed Acyclic Graph (DAG) representing our study hypothesis on the interconnected relationship between motor and sleep disturbances with PVS alterations and cognition in Parkinson’s disease, including the covariates used in this study.

The objective of this cross-sectional study is to investigate the relationship between PVS, sleep quality, and walking difficulties in a cohort composed of healthy controls, prodromal PD individuals, and patients with a clinical PD diagnosis. We *hypothesize* that individuals exhibiting walking difficulties may also experience poorer sleep quality, affecting brain waste clearance via the glymphatic system and ultimately cognition.

We first assessed the association between PVS and walking difficulties, as well as the link between walking difficulties and sleep problems, accounting for education, age, sex, study group, magnetic field strength and body mass index (BMI)^14^. This analysis was repeated including APOE4 genotype as a covariate, to assess the effect of this genetic information on the relationship between PVS and walking difficulties. We further analyzed the association between sleep problems and PVS, accounting for the same covariates listed above.

Given our interest in the role of PVS in cognition, particularly in the context of walking and sleep disturbances, we aimed to investigate whether motor and sleep disturbances could affect glymphatic function, quantified by PVS volume fraction, and ultimately impact cognition. We hypothesized that PVS mediates the relationship between walking difficulties, higher level of sleep deficits and cognitive performance. Motor dysfunction has been linked to larger PVS in PD, leading to impaired cerebral blood flow and increased oxidative stress, previously associated with cognitive decline^15,16^. As such, walking difficulties may be an early marker of microvascular pathology, with PVS enlargement serving as a biological intermediary linking motor decline to impaired cognition. Poor sleep similarly disrupts waste clearance and has been tied to PVS enlargement^5^, which may promote neuroinflammation in regions critical for both movement and cognition^17^.

We finally investigated whether PVS volume fraction could predict PD disease severity, measured by both biological markers, such as CSF alpha-synuclein at baseline, longitudinal beta-amyloid, t-Tau and phosphorylated Tau (p-Tau), and cognitive scores after five years. CSF biomarkers have been previously considered as markers for progression of cognitive decline in PD^13^, as well as associated with worsening of motor symptoms^18^.

## 3. Results

Participants divided into HC, prodromal and PD patients differed in education, walking difficulties and magnetic field strength (Table 1, *p-values*<.001) and SDM scores (*p*=.002). Groups did not differ in insomnia levels (*p*=.4).

**Table1:**
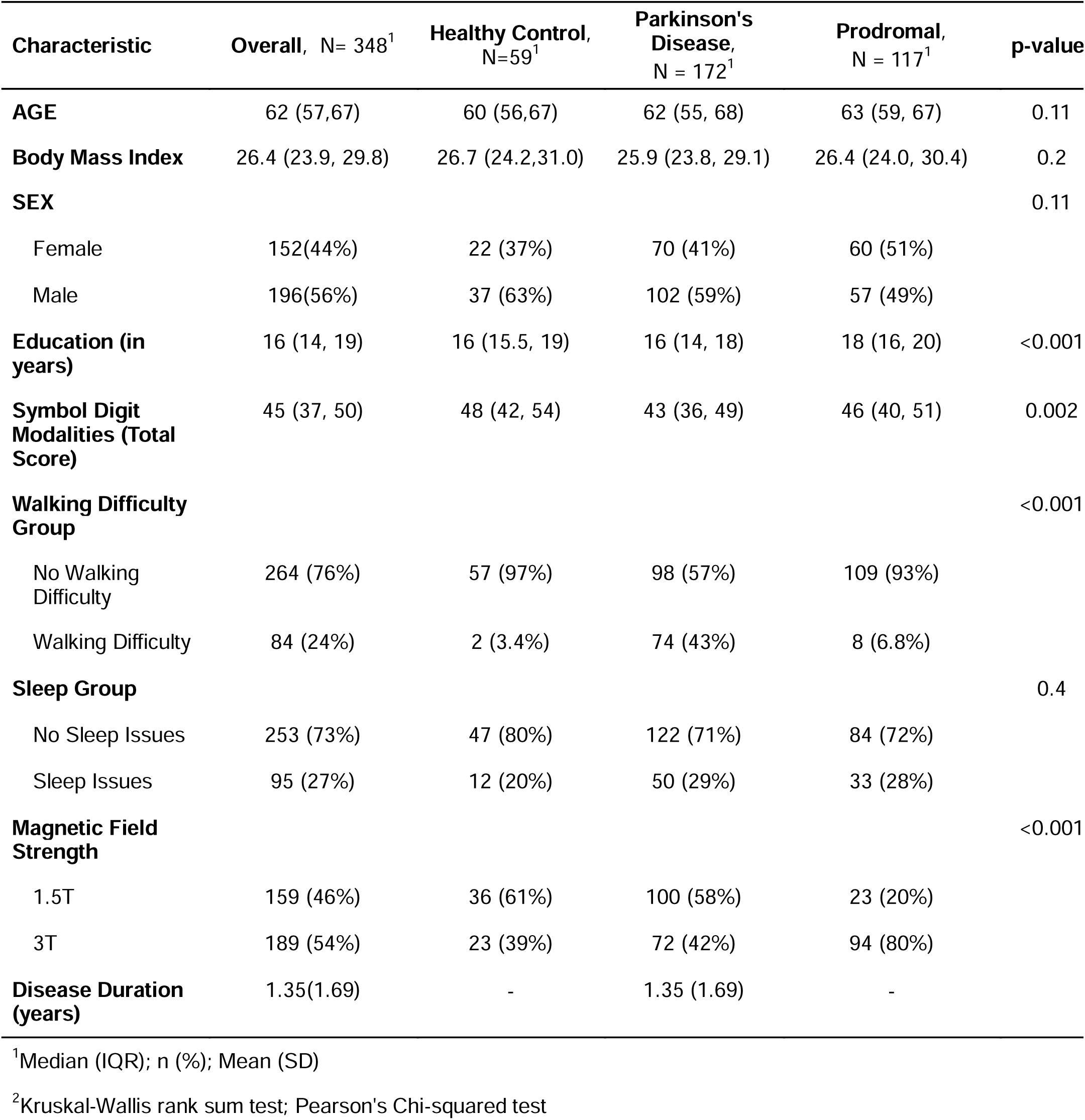
Demographic information of this study sample. Walking difficulties were assessed with Unified Parkinson’s Disease Rating Scale (MDS-UPDRS) II (Item 2.12). Sleep problems were assessed based on the insomnia levels from the MDS-UPDRS I (Item 1.7).

### 3.1. Association of PVS with walking difficulties and sleep problems

The GLM showed no statistically significant relationship between WM-PVS volume fraction and walking difficulties (beta(standard error(SE)) =.12(.08), *p*=.12), with or without APOE4 in the model as a covariate (beta(SE)=.13(.079), *p*=.10). We also found no statistically significant association between BG-PVS and walking deficits (beta(SE)=.016(.12), *p*=.89), nor when considering APOE4 as a covariate (beta(SE)=.019(.11), *p*=.87). Results of the logistic regression showed no significant associations between PVS and sleep problems, neither in WM-PVS (beta(SE)=- .05(.06), *p*=.45) nor in BG-PVS (beta(SE)=-.12(.13), *p*=.23). We additionally examined the link between walking difficulties (dependent variable) and sleep problems through a logistic regression, adjusting for age, sex, education, study group, magnetic field strength and BMI; we observed a significant positive association (beta estimate=0.82, SE=0.32, *p*=0.0099).

There was a significant effect of study group for the PD patient group on walking difficulties (**Figure3**).

**Figure2:**
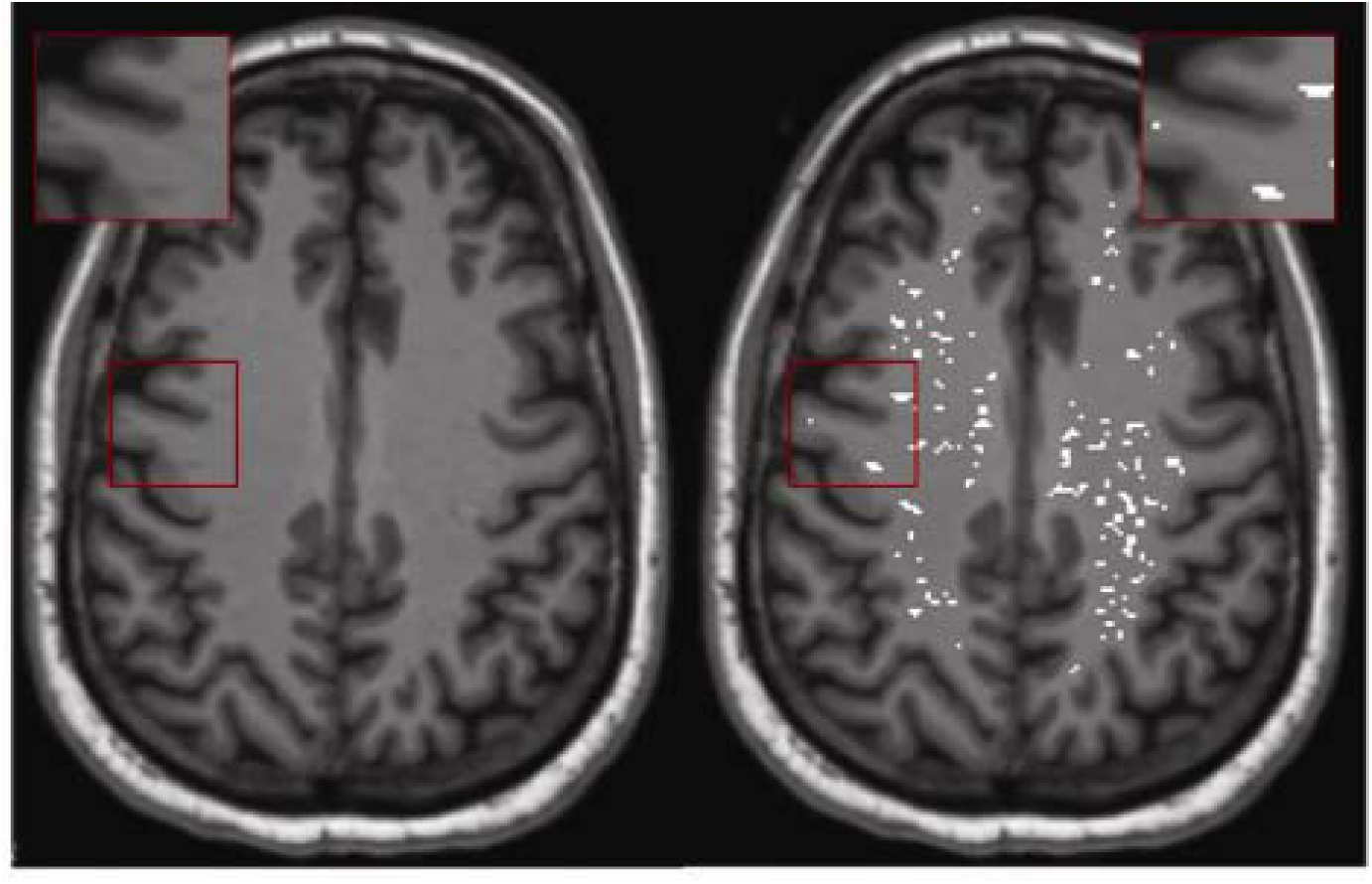
Example of PVS segmentation result in one participant.

**Figure3:**
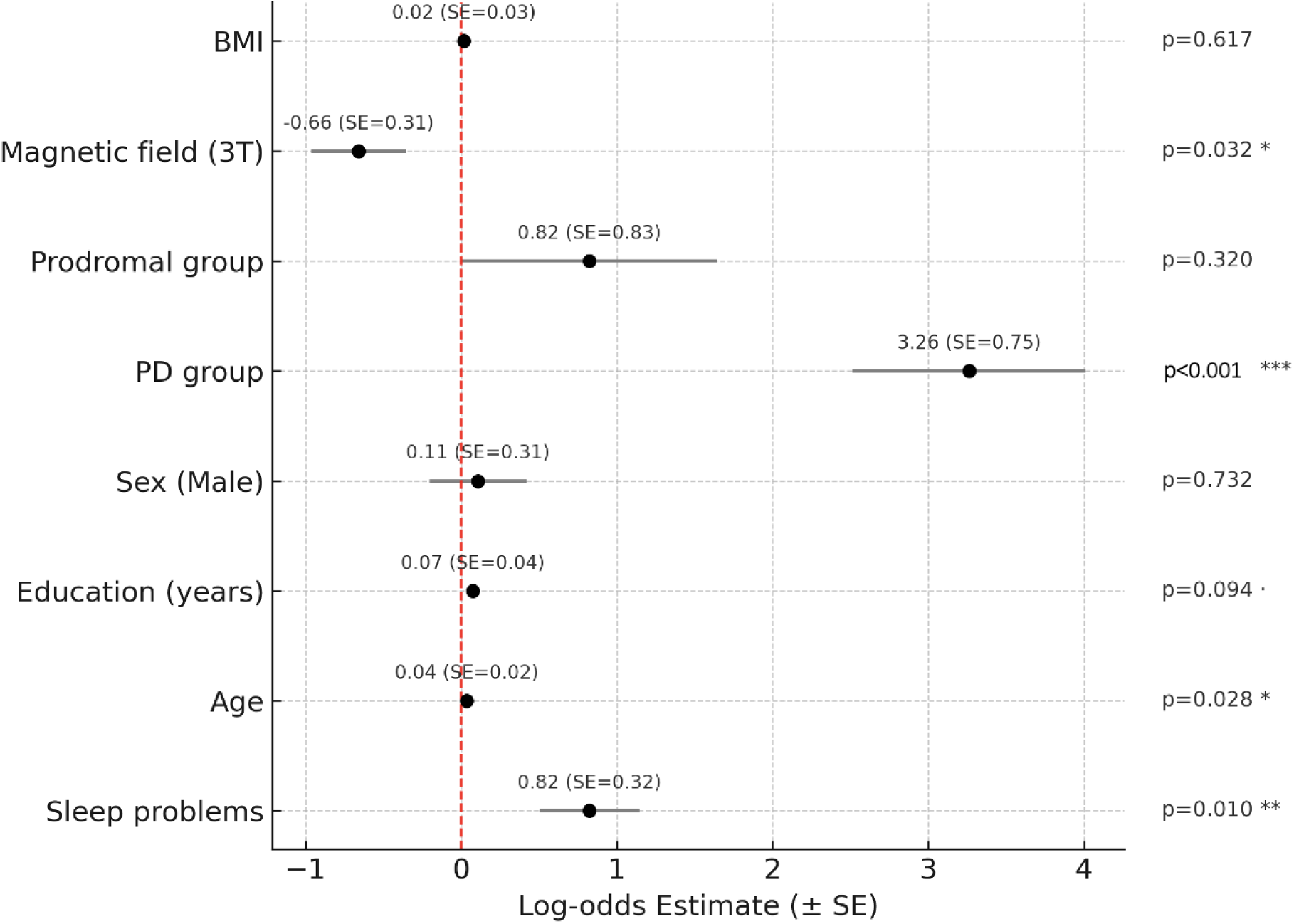
Results of the logistic regression used to investigate the link between walking difficulties (binary dependent variable) and sleep problems. The walking difficulties variable was dummy coded as 0=people who had walking score=0 and 1=people with any level of walking difficulties – walking score=1,2,3,4.

To further explore our findings, we repeated the analyses with regional PVS based on a WM atlas–based parcellation; however, no significant associations were found after correction for multiple comparisons.

### 3.2. SEM analysis: walking and sleep - PVS - cognition

The SEM analysis revealed both direct and total effects of walking behavior on cognition. Walking difficulty showed a significant direct effect on SDM performance (estimate=−.288, SE=.11, 95%CI [−.507,−.070], *p*=.01) (**Figure4a**). The effect of walking difficulty on WM-PVS was positive but non-significant (estimate=.233, SE=.15, 95%CI [−.064,.528], *p*=.119). The indirect effect of walking on cognition through WM-PVS was not significant (estimate=-.011, SE=.014, 95%CI [−.013,.02], *p=*.4).

**Figure4:**
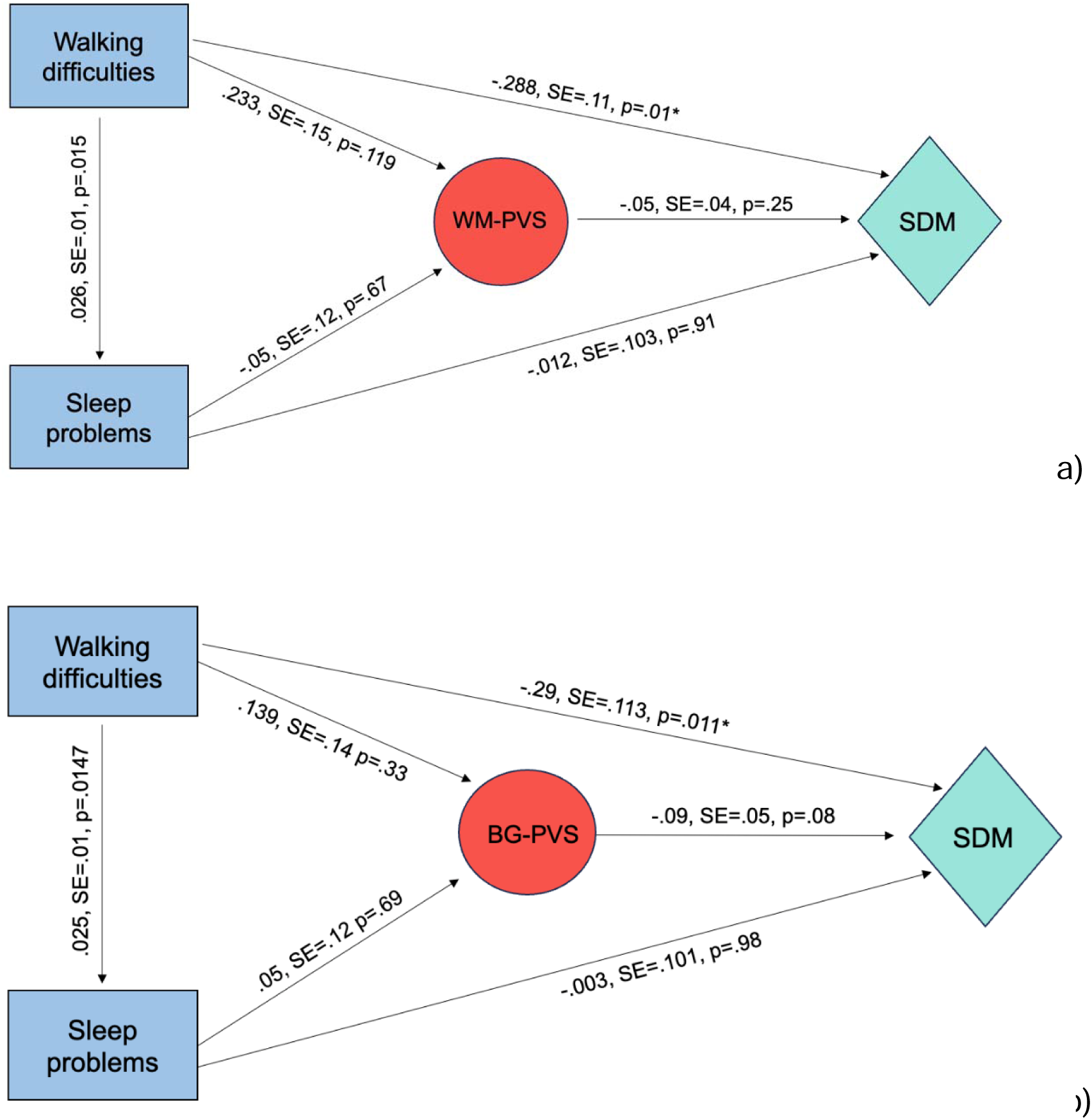
SEM analysis testing the relationship between walking difficulties and cognition, with a) WM-PVS and b) BG-PVS as mediator.

For sleep, the direct effect on SDM performance was small and non-significant (estimate=−.012, SE=.103, 95% CI [−.214,.192], *p*=.91), and the effect of sleeping difficulties on PVS was not significant (estimate=-.05, SE= .12, 95% CI [−.28, .19], *p*=.67). The indirect effect was not significant (estimate=.002, SE=.008, 95%CI [− .01,.02], *p=*.75). Consistent with the logistic regression analysis reported above, the correlation path between walking deficits and sleep problems showed a significant positive association (estimate=.026, SE=.01, 95%CI [.005, .04], *p*=.015). The SEM model yielded the following model fit measures: chi-square (χ²)= 94.7, p < .001, comparative fit index (CFI) = 0.72, Root Mean Square Error of Approximation (RMSEA) = 0.129, Standardized Root Mean Square Residual (SRMR) = 0.07.

When BG-PVS was tested as the mediator, walking difficulties remained significantly related to cognition. The total and direct effect on SDM performance was negative and significant (estimate=−.29, SE=.113, 95%CI [−.508,−.07], *p*=.011). The path from walking to BG-PVS itself was small and non-significant (estimate=.139, SE=.14, 95%CI [−.123,.42], *p*=.31). The path from BG-PVS to cognition trended toward significance (estimate=−.09, SE=.05, 95%CI [−.198,−.002], *p*=.08) (**Figure4b**). The indirect effect was negative but not significant (estimate=-.012, SE=.016, 95%CI [−.05,-.012], *p*=.42). For sleep, no meaningful associations with cognition were observed. The direct effect on SDM performance was negligible (estimate=−.003, SE=.101, 95%CI [−.207,.193], *p*=.98). Likewise, the relationship between sleep and PVS was not significant (estimate=.05, SE =.12, 95%CI [−.18,.28], *p*=.69). The indirect effect of sleep on cognition through BG-PVS was not significant (estimate=-004, SE=.01, 95%CI [− .02,.02], *p=*.71). The correlation path between walking deficits and sleep problems showed a significant positive association (estimate=.027, SE=.01, 95%CI [.005, .05], *p*=.0149).

Fit statistics for the SEM model were as follows: χ²= 93.06, p<.001, CFI=.77, RMSEA=.13, SRMR=.07.

We repeated the same analyses within the PD patients to examine differences in PVS relationship with walking difficulties and sleep problems in the clinical subgroup. Within-group results confirmed our findings in the whole sample, showing a significant direct association between walking deficits and cognitive levels, not mediated by PVS. Detailed results are reported in **Supplementary Results**.

### 3.3. Baseline PVS and disease progression

The longitudinal analysis indicated no statistically significant predictive effects of baseline PVS on CSF biomarkers (n=132) at 5 years. With t-Tau, this was seen both in WM-PVS (beta(SE)=84.09(301.74), *p*=.78) and BG-PVS (beta(SE)=.-85.11(846.88), *p*=.92). Similarly, analyses involving p-Tau and beta-amyloid revealed no significant predictions.

We examined the relationship between WM-PVS and BG-PVS burden and baseline alpha-synuclein (n=141). No significant associations were observed. Finally, we looked at cognition score 5 years later. While WM-PVS did not show any significant predictive effect on 5-year SDM scores, BG-PVS was significantly predictive of cognitive function 5 years later (estimate=-308.8, SE=113.4, *p*=.007), alongside other risk factors such as older age, male sex, walking difficulties, and Parkinson’s disease, while higher education offered protection.

## 4. Discussion

Utilizing the PPMI cohort data, we found that PVS does not mediate the relationship between walking difficulties, sleep problems and cognition. Our results demonstrated a direct relationship between the gait symptoms and cognitive performance, as well as between sleep disturbances and walking deficits. Finally, the longitudinal analysis revealed that baseline PVS does not predict CSF protein markers accumulation, but BG-PVS predicts cognitive status 5 years later.

Enlarged PVS may reflect impaired brain clearance, leading to CSF buildup of waste molecules. PVS enlargement has been previously associated with cerebro-vascular diseases and other conditions in older adults, such as depression, diabetes, and neurodegenerative diseases^34^. Both global and regional PVS volumes were found to be significantly larger in individuals with PD compared to HC^35^. Alongside our results, finding a negative association between BG-PVS and cognitive scores measured 5 years later, these observations suggest that PVS changes could influence various neurodegenerative pathology and act as early markers for cognitive decline.

Our results showed PVS does not mediate the observed relationship between walking problems, sleep disturbances and cognition.

Others have reported the opposite results, finding that baseline PVS was related to longitudinal motor dysfunction in PD, but not cognitive decline^36^. Mixed results have been reported on the association between PVS and cognition. Previous studies showed a negative relationship between PVS volume fraction and cognition in patients with early-stage PD^35^, whereas others found no association in patients with ischemic stroke^16^ or PD^36^. It is possible our model could not detect such a relationship due to the variables chosen to represent gait and sleep difficulties.

In our analyses, the relationship between walking difficulties and PVS burden and sleep problems and PVS, was not significant. This lack of an initial association indicated that mediation analysis was not methodologically justified; we instead applied a structural equation modeling framework to assess the interrelationships among the three variables within a single model.

Our SEM mediation analyses on BG-PVS and WM-PVS revealed that severe walking deficits are directly related to SDM lower scores, and PVS did not mediate such effects, opposite to our hypothesis.

The observed direct relationship between walking difficulties and cognitive performance in our cohort builds on previous studies linking motor and cognitive decline in PD, especially in attention and frontal-executive function^15^. A robust body of literature shows gait problems contribute to and/or co-occur with cognitive decline in PD. Greater motor impairment was linked to lower cognitive scores, with MDS-UPDRS balance-gait measure found to be the best predictor of cognitive impairment and dementia^37^. Walking disturbances have been previously recognized as a marker of cognitive dysfunction even in prodromal stages of neurodegenerative diseases, and gait performance worsened as cognitive impairment progressed^15^. Mild cognitive impairment and dementia were also previously associated with worsening of non-motor symptoms, including excessive sleepiness^38^; some studies have found that patients with REM-sleep behavior disorder have a higher rate of cognitive decline^39^. The directional link between motor dysfunction and cognitive decline is also supported by a previous study showing that the reverse association is not significant, such that cognitive reserve (the brain’s ability to compensate for decline) did not influence gait performance in patients with idiopathic PD^40^.

Our findings indicate that sleep problems are associated with greater walking difficulties. While sleep disturbances contribute directly to motor dysfunction^41^, the underlying mechanisms may not primarily involve cerebrovascular changes, specifically PVS dilation. Previous studies on PD reported how individuals with higher levels of insomnia showed worse gait performance and increased risk of falls^42^; the presence and level of REM sleep without atonia (RSWA) was also related to the severity of gait disturbances in people with PD^43^. Disrupted sleep patterns have been reported to impair motor function; it is proposed that sleep restorative effects are compromised for regions involved in motor control, such as the basal ganglia^7^.

Cognitive decline in PD often manifests as impairment of processing speed, captured by SDM. Slower motor processing and execution of motor tasks is thought to be driven by dopaminergic deficits in the basal ganglia, which also contribute to slower processing speed and attention deficits^44^. The negative association between baseline BG-PVS and SDM performance at both baseline and follow-up suggests that cognitive decline in PD is linked to basal ganglia pathology, rather than global PVS burden. Unlike WM-PVS, BG-PVS appear more relevant for detecting early cognitive changes and tracking disease progression. This aligns with prior work showing that enlarged BG-PVS predicted longitudinal decline in processing speed, executive function, and episodic memory over five years, implicating disruption of frontal–subcortical circuits^45^.

Previous literature suggests that CSF biomarkers reflect disease progression in the prodromal and early stages of PD^41^. At these stages, alterations in CSF alpha-synuclein, amyloid-beta, and Tau may signal early neurodegeneration before major motor symptoms appear. Mollenhauer and colleagues found reduced CSF alpha-synuclein levels were associated with more severe motor symptoms and predicted disease worsening in the early phases of PD^46^. Increased levels of p-Tau and decreased levels of Aβ42 were previously linked to PD cognitive impairment and disease progression^47^; conversely, when looking at the relationship between PVS and CSF biomarkers change, neither we nor other reports from PPMI data have shown an association between PVS volume fraction and longitudinal change in CSF biomarkers^48^. One possible explanation to our negative finding is that the patients with PD in the PPMI dataset were very mildly affected when recruited, by definition not yet on medications for PD. The first few years of PD progression tend to be quite mild and generally does not include significant cognitive impairment or disability.

One limitation of our study is the lack of alpha-synuclein longitudinal data five years after the initial measurements, preventing a clearer understanding of disease progression and symptom development. Another important consideration is the study’s primarily cross-sectional design, which limits our ability to determine the temporal directionality of the associations observed between walking difficulties, sleep disturbances, and PVS enlargement. It remains unclear whether PVS enlargement contributes to motor and sleep impairments, or whether gait and sleep dysfunction influence PVS structure. Additionally, in our analysis of APOE4 genotype, participants were categorized based on the presence or absence of at least one APOE4 allele. Further stratification by number of APOE4 alleles was not performed due to the small number of individuals carrying two APOE4 allele copies. Despite these limitations, our study offers novel insights and several strengths. We utilized a semi-automated segmentation method for PVS identification, compared to previous studies that relied on manual counting. Importantly, we further excluded WMH as potential false positives, ensuring a more accurate assessment of PVS volume fraction. Future work should consider additional PVS structural features, such as tortuosity, shape, and size, which may offer more nuanced insights into the clinical relationships between PVS, sleep disturbances, and motor dysfunction in PD. Examining sleep and gait disturbances longitudinally will also be important for understanding how PVS enlargement may influence sleep quality and motor deficits over time and for untangling temporal relationships among walking and sleep impairments. The SEM approach enabled us to test the hypothesized associations specified a priori. While certain fit indices did not meet conventional thresholds, the observed associations were consistent with previous research and support the reliability of the findings. Future research should incorporate additional variables and longitudinal designs to further refine our understanding of the mechanisms linking walking, sleep, PVS burden, and cognition. In PD, the integration of behavioral and clinical factors, such as walking difficulties, sleep disturbances, genetics, clinical diagnosis and biological markers, with neuroimaging markers like PVS adds novelty to our study, as previous studies considered these aspects separately. This multidimensional approach not only deepens our understanding of Parkinson’s disease pathophysiology, but highlights the importance of integrating clinical, biological, and structural factors to capture the complexity of disease progression, potentially guiding future therapeutic strategies and personalized care.

## 2. Methods

### 2.1. Study participants

Participants were from the Parkinson’s Progression Markers Initiative (PPMI)^19^. PPMI is a comprehensive international study launched in 2010 to identify and validate biomarkers for PD progression. The sample used in this study is from the PPMI Clinical, which includes in-person clinical and imaging assessment at fifty multi-international sites. Data used in the preparation of this article was obtained on 2023-07-18 from the Parkinson’s

Progression Markers Initiative (PPMI) database (www.ppmi-info.org/access-dataspecimens/download-data), RRID:SCR_006431. For up-to-date information on the study, visit www.ppmi-info.org. We used participants’ baseline data, with an initial sample size of *N*=431. We excluded participants who had missing neuroimaging data (structural T1w images and PVS segmentation output) and missing data in demographics (age, sex, study group, BMI, education), walking difficulties and sleep scores, resulting in a sample of n=365. Of the 365 participants, 11 participants failed Freesurfer^20^ preprocessing. An additional 6 were missing magnetic field scanner information. These 17 participants were excluded, and the pipeline for segmenting the PVS was applied to the remaining 348 participants. The last step of our pipeline removed white matter hyperintensities (WMH), often registered as PVS false positives.

Our final sample size was *N*=348, (age range 31-82 years old), including 172 PD patients, 117 prodromal cases and 59 HC. Participants were categorized as PD patients if they demonstrated a combination of bradykinesia and resting tremor and/or rigidity within two years of diagnosis. Prodromal PD cases were defined by olfactory loss or RBD with dopamine transporter (DAT) deficit^19^. HC subjects were clinically ascertained as not having any neurologic dysfunction and Montreal Cognitive Assessment (MoCA)>26. Demographic information is reported in **Table1**.

### 2.2. Neuropsychological and motor variables

Walking difficulties were assessed with the Movement Disorder Society-Unified Parkinson’s Disease Rating Scale (MDS-UPDRS) II (Item 2.12). The MDS-UPDRS is divided into four sections; the first and second parts evaluate non-motor and motor experiences of daily living respectively, the third part focuses on motor examination, and the fourth part assesses motor complications. Item 2.12 assesses level of walking and balance difficulties, represented by a Likert scale, with higher scores reflecting more difficulties (0=normal, 1=slight, 2=mild, 3=moderate, 4=severe). The population was sub-divided by dummy coding the walking difficulties variable (0=people who had walking score=0, 1=people with any level of walking difficulties – walking score=1,2,3,4) (**Table1**, variable named ‘Walking difficulty group’).

Sleep problems were assessed from insomnia ratings from the MDS-UPDRS I (Item 1.7), evaluating trouble staying asleep on a scale from 0 to 4, with higher scores indicating worse insomnia. In our analysis, the sleep variable was dummy coded, based on a cut-off value from a previous study^21^: individuals with sleep scores of 0 or 1 were assigned a value of 0, while those with a sleep score of 2 or higher were coded as 1, thus binarizing our participants as good and poor sleepers.

Cognition was assessed for each participant with the total score of Symbol Digit Modalities (SDM) test, a neuropsychological assessment that measures processing speed, attention, and visual-motor coordination. Participants are given a key that pairs specific symbols with numbers. They must then quickly match symbols to their corresponding numbers. The score is the number of correct symbol-number pairings made within 90 seconds. Higher scores indicate better cognitive functioning and faster processing speed.

We chose SDM as the cognitive measure, as it was previously shown in PPMI data that SDM performs better than other cognitive scales in detecting cognitive impairment and progression in PD^22^, based on its steady downward trend, superior reliability, enhanced capability in detecting MCI, and significant association with motor progression, both at baseline and longitudinally.

Average scores for the total sample and each subgroup are found in **Table1**. The PD group showed the lowest SDM median and interquartiles compared to prodromal and HC groups.

### 2.3. Imaging

Imaging data were acquired according to the PPMI Clinical MRI Technical Operations Manual version 7.0 (2015) described previously^23^. 3D T1-weighted images at baseline were acquired using 1.5 and 3 Tesla scanners, with a slice thickness of 1.5 mm or less with no interslice gap; voxel size was 1×1x1.2mm. A high-resolution 3D T2 FLAIR sequence was also acquired, with a slice thickness of 1.2 mm, voxel size 1×1 mm in plane resolution, with a total scan time between 20-30 minutes. Additional information can be found in the MRI Technical Operations Manual, Final version 7.0 (2015).

Freesurfer (v.5.3) was used for image preprocessing, which included motion correction, image normalization, and skull stripping. Brain volume and parcellation were obtained with the recon-all module on the Laboratory of Neuro Imaging (LONI) pipeline system (https://pipeline.loni.usc.edu)^24,25^.

PVS were mapped and segmented from T1w images, following a previously published technique^26,27^. A non-local mean filtering method was applied to denoise the MRI T1w images^28^, which defines the similar intensity of a voxel to its neighboring voxels. A Frangi filter was further applied at a voxel level^29^ using the Quantitative Imaging Toolkit (QIT)^30^, to detect tubular PVS structures on the T1w, and obtaining a probabilistic map of vesselness^29^. To capture true positives and true negatives, a threshold of 0.00003 in WM-PVS was applied to obtain a binary PVS mask for data acquired at 3T, and a threshold of 0.0003 to data acquired at 1.5T. For BG-PVS the threshold of 0.0004 was applied for data acquired at 1.5T and threshold of 0.0001 for 3T data. An example segmentation result is provided in **Figure2**. The WM-PVS volume fraction was calculated by dividing the total WM-PVS volume by the total white matter volume for each participant. The BG-PVS volume fraction was calculated by dividing the total BG-PVS volume by the basal ganglia volume. For regional PVS evaluation, the PVS volume fraction was determined by dividing the PVS volume within each regional white matter area by the corresponding total white matter volume. This calculation was performed across all WM regions defined by the Desikan–Killiany atlas.

In PPMI, FLAIR data is available, allowing subtraction of WMH as false positives from the PVS segmentation mask. The mask for WMHs was created by combining the WMH masks generated with the Lesion Growth Algorithm^25^, which measures the intensity distribution in white matter, grey matter, and CSF on FLAIR and T1-weighted images and identifies WMH voxels as outliers in the signal distribution curve, with neighboring voxels iteratively assigned to the WMH mask by weighing the likelihood of belonging or not to the WMH mask, and the Sequence Adaptive Multimodal Segmentation tool^24^, a contrast-adaptive method based on a generative approach in which a forward probabilistic model is inverted to obtain automated and robust segmentations of WMH. Voxels marked as WMH were finally subtracted from the PVS mask to remove false positive PVS^24,25^.

### 2.4. CSF biomarkers

CSF samples, collected according to the PPMI Laboratory Manual procedure, were utilized to measure alpha-synuclein, amyloid-beta, t-Tau and Tau phosphorylated at the threonine-181-position (p-tau181), using Elecsys electro-chemi-luminescence immunoassays on the cobase 601 platform (Roche Diagnostics)^31,32^ as biomarkers of PD progression^13^. While amyloid-beta, t-Tau and p-Tau were collected again after 5 years, alpha-synuclein was not measured at that timepoint. As a result, analyses involving alpha-synuclein and PVS were limited to baseline cross-sectional data. The alpha-synuclein assay has a measurement range of 300-7000 pg/mL, the amyloid-beta assay of 200 to 1700 pg/mL, the t-tau assay: 80 to 1300 pg/mL, and the p-tau181 assay: 8 to 120 pg/mL.

### 2.5. Statistical analysis

The initial analysis examined the association between walking difficulties and PVS volume fraction (dependent variable) with BMI, age, sex, education, magnetic field strength, and study group as covariates. Age, BMI and education were modeled as continuous variables. We used a generalized linear model (GLM) with Gamma distribution and log link to consider the positive skewed distribution of PVS volume fraction. The association between the binary sleep variable and PVS was assessed using logistic regression with a binomial family and a logit link function.

We utilized APOE4 genotype data to consider possible confounding on the relationship between PVS and walking difficulties. The APOE4 genotype was added to the model above as a covariate. The genotype information was dummy coded as 0/1, with 0=‘absence of APOE4 allele’ and 1=’presence of APOE4 allele’. Genetic information was missing for one participant.

We employed structural equation modeling (SEM)^33^ to evaluate the direct and indirect effects of walking and sleep on cognitive performance, with PVS measures as mediators. Because WM-PVS and BG-PVS are considered to reflect distinct underlying vascular mechanisms, we modeled them as separate mediators in independent SEM analyses. Cognitive performance, measured by SDM, was specified as the outcome. Walking and sleep disturbances were modeled as exogenous binary predictors, and a correlation path between them was included. Age, years of education, sex, prodromal cohort status, PD cohort status, magnetic field strength, and BMI were included as covariates in all paths with PVS or cognition as outcomes. Indirect effects of walking and sleep on cognition via WM-PVS and BG-PVS were computed along with the direct and total effects. Standard errors (SE) and confidence intervals (CIs) for indirect effects were derived using bootstrap resampling to account for non-normal sampling distributions. Study group was modeled using two indicator variables, with healthy controls serving as the reference category.

Thus, coefficients for these indicators reflect differences in the outcome variable between prodromal or PD participants and healthy controls. SEM analyses were also repeated within the PD patients’ group only.

One of our aims was to explore whether the PVS volume fraction could predict CSF biomarkers accumulation, including t-Tau, p-Tau, and amyloid-beta, after a 5-year period. We analyzed data from 132 participants (44 HC, 7 prodromal, 81 PD) who had longitudinal CSF biomarker information available. We also investigated whether the PVS volume fraction at baseline could predict future cognitive scores (longitudinal SDM scores were available for n=275, 57 HC, 138 PD, 80 prodromal). We used linear regression models for all analyses, with 5-year SDM or CSF biomarker values as the dependent variable and PVS as independent variables. Covariates were age, sex, study group, education, walking difficulties, sleep, magnetic field strength and BMI. We conducted sensitivity analyses by fitting the same models without adjusting for scanner. Statistical significance corresponded to 2-sided *p*-value<0.05.

## Supporting information

Supplementary Results

## Data Availability

The data used in this study are available from the Parkinson’s Progression Markers Initiative (PPMI) database (https://www.ppmi-info.org/). Access to the PPMI data requires registration and approval from the PPMI Data and Publications Committee. All researchers may apply for access via the PPMI website.

## Acknowledgments

This study was funded by National Institutes of Health (NIH) grant number R01AG070825. The funder played no role in study design, data collection, analysis and interpretation of data, or the writing of this manuscript.

## Authors’ contribution

**Conceptualization:** Francesca Sibilia.

**Data curation:** Francesca Sibilia, Giuseppe Barisano, Maryam Fotouhi, Jeiran Choupan.

**Formal analysis:** Francesca Sibilia, Wendy J. Mack.

**Funding acquisition:** Jeiran Choupan.

**Investigation:** Francesca Sibilia.

**Methodology:** Francesca Sibilia, Giuseppe Barisano, Jeiran Choupan.

**Project administration:** Francesca Sibilia.

**Resources:** Jeiran Choupan, Giuseppe Barisano.

**Software:** Jeiran Choupan, Giuseppe Barisano.

**Supervision:** Jeiran Choupan.

**Validation:** Francesca Sibilia.

**Visualization:** Francesca Sibilia.

**Writing-original draft:** Francesca Sibilia.

**Writing-review & editing:** Francesca Sibilia, Giuseppe Barisano, Maryam Fotouhi, Philipp A. Loehrer, Katherine Wong, Wendy J. Mack, Mark Lew, Haidar Dafsari, Jeiran Choupan

## Disclosures and Competing Interests

Philipp A. Loehrer was funded by the von Behring-Röntgen-Foundation and a research grant of the University Medical Center Giessen and Marburg (UKGM). The perivascular space mapping technology is part of a patent number US11908131B2 owned by JC. JC declares status as employee of the private company NeuroScope Inc (NY, USA). All other authors declare no financial or non-financia competing interests.

### Ethical standard

All procedures performed in studies involving human participants were in accordance with the ethical standards of the institutional and/or national research committee and with the 1975 Helsinki declaration and its later amendments or comparable ethical standards. Informed consent was obtained from all the patients included in the study.

