## Supplementary Results for "Independent Associations of Sleep and Walking with Cognition in Parkinson’s Disease: A Perivascular Spaces Analysis"

*1. Mediation analyses in each clinical group*

We stratified our population based on the clinical diagnosis in healthy controls (n=59), prodromal cases (n=117) and PD patients (n=172). We replicated the same analyses protocols as for the whole population within the PD group. Here we report the main significant results.

*1.1. Patients with Parkinson's disease*

The SEM analysis revealed both direct and total effects of walking behavior on cognition. Walking showed a significant direct effect on SDM performance (estimate=−.303, SE=.12, 95% CI [−.544, −.065], p=.012). Although the path from walking to WM-PVS was positive but non-significant, the total effect of walking on cognition was also significant (estimate=−0.317, SE=.122, 95% CI [−.561, −.07], p=.009). This indicates that walking exerted a reliable overall influence on cognitive performance when both direct and indirect pathways were considered. For sleep, neither the direct nor total effects reached significance. The model showed the same trends when BG-PVS was considered as mediator.

*2. Does sleep mediate the relationship between walking difficulties and PVS?*

Another mediation analysis was conducted to examine whether sleep mediates the relationship between walking difficulties and PVS. The mediation analysis did not reveal a significant mediating effect of sleep in the relationship between walking difficulties on PVS, although a significant positive association was found between sleep and walking deficits (estimate = .64, $p$ =.048). We found no significant results when running this model with the basal ganglia as mediator.

*3. Mediation analyses with MoCA as cognitive measure*

To assess the robustness of our findings on cognition, we repeated the mediation analysis using the Montreal Cognitive Assessment (MoCA) total score as an alternative cognitive measure. The MoCA is a widely used global cognitive screening tool in PD, capturing a broader range of cognitive domains than the SDM. This analysis also did not reveal significant indirect effects through PVS nor direct effect between walking deficits and MOCA scores, suggesting that the association between walking difficulties, PVS burden, and cognition may involve domain-specific processes that are not fully captured by a global cognitive measure.
